# InSilicoSeq 2.0: Simulating realistic amplicon-based sequence reads

**DOI:** 10.1101/2024.02.16.580469

**Authors:** Stefan H. Lelieveld, Thijs Maas, Tessa C. X. Duk, Hadrien Gourlé, Henk-Jan van den Ham

## Abstract

**Motivation:** Simulating high-throughput sequencing reads that mimic empirical sequence data is of major importance for designing and validating sequencing experiments, as well as for benchmarking bioinformatic workflows and tools.

**Results:** Here, we present InSilicoSeq 2.0, a software package that can simulate realistic Illumina-like sequencing reads for a variety of sequencing machines and assay types. InSilicoSeq now supports amplicon-based sequencing and comes with premade error models of various quality levels for Illumina MiSeq, HiSeq, NovaSeq and NextSeq platforms. It provides the flexibility to generate custom error models for any short-read sequencing platform from a BAM-file. We demonstrated the novel amplicon sequencing algorithm by simulating Adaptive Immune Receptor Repertoire (AIRR) reads. Our benchmark revealed that the simulated reads by InSilicoSeq 2.0 closely resemble the Phred-scores of actual Illumina MiSeq, HiSeq, NovaSeq and NextSeq sequencing data. InSilicoSeq 2.0 generated 15 million amplicon based paired-end reads in under an hour at a total cost of €4.3e^-05^ per million bases advocating for testing experimental designs through simulations prior to actual sequencing.

**Availability and implementation:** InSilicoSeq 2.0 is implemented in Python and is freely available under the MIT licence at https://github.com/HadrienG/InSilicoSeq

## Introduction

Amplicon-based sequencing is considered the *de facto* standard in the characterization of specific target regions of a genome. Examples are the targeted sequencing of genetic markers such as the 16S rRNA gene for bacteria^1^, the internal transcribed spacer (ITS) region for fungi^2^, and the repertoire of the adaptive immune receptor repertoire (AIRR) of B and T cells^3,4^. However, generating experimental sequencing data can be time-consuming and resource intensive. Therefore, *in silico* simulations of amplicon sequenced data are an important tool to test and validate sequencing experiments as well as to benchmark downstream workflows and tools.

Simulating amplicon-based next generation sequencing (NGS) experiments can be utilized to test a variety of parameters to optimize sequencing strategies as well as the performance of bioinformatics tools and pipelines^5^. This includes evaluating the effects of varying sequencing depth, assessing the impact of different error rates, and studying the impact of sequencing technologies^6^. Thus, the insights gained from running simulated data guides the scientist to make an informed decision on what approach will give the desired results.

Here, we present an updated and extended version of InSilicoSeq, v2.0, a Python-based read simulation package that contains many new features, including amplicon-based sequencing and new error models for the latest Illumina sequencing platforms. Existing features have been improved resulting in a constant memory-footprint, corrected read-sampling and fragment lengths, high-resolution logging of the sequencing errors, and reads that closely resemble the Phred-scores of real reads. We demonstrate the novel and easy-to-use amplicon sequencing algorithm of InSilicoSeq 2.0 by AIRR sequencing data of paired-end 301 base-pair long reads. The software-package is freely available on https://github.com/HadrienG/InSilicoSeq.

## 2. Features and benchmarks

### 2.1 Features

As described in the original publication, InSilicoSeq can simulate metagenomic data; it can accurately model Phred^7^ scores based on real sequencing data; it supports substitution, insertion, and deletion errors; and it supports insert size distribution and GC bias by implementing Kernel Density Estimation (KDE)^8^. In line with simulating amplicon-based sequencing data (Supplementary figure 1), new error models based on the Illumina MiSeq and NextSeq sequencing platforms (paired-end reads of 301bp) have been included in the software package (See supplementary table 1 and supplementary figure 2 for a complete overview). For the MiSeq platform, a total of five error models have been generated each based on reads with a different average Phred-score (20, 24, 28, 32, and 36; figure 1F-H and supplementary figure 2). This allows simulating sequencing reads with a wide variety of qualities. The NextSeq reads showed high quality scores and therefore only a single error model was constructed for this platform (Figure 1D-E). The MiSeq error models and the NextSeq model are based on the data of all eight SRA entries in project PRJEB20178^9^ and Illumina base space demo data project 370456276^10^, respectively.

**Figure 1.**
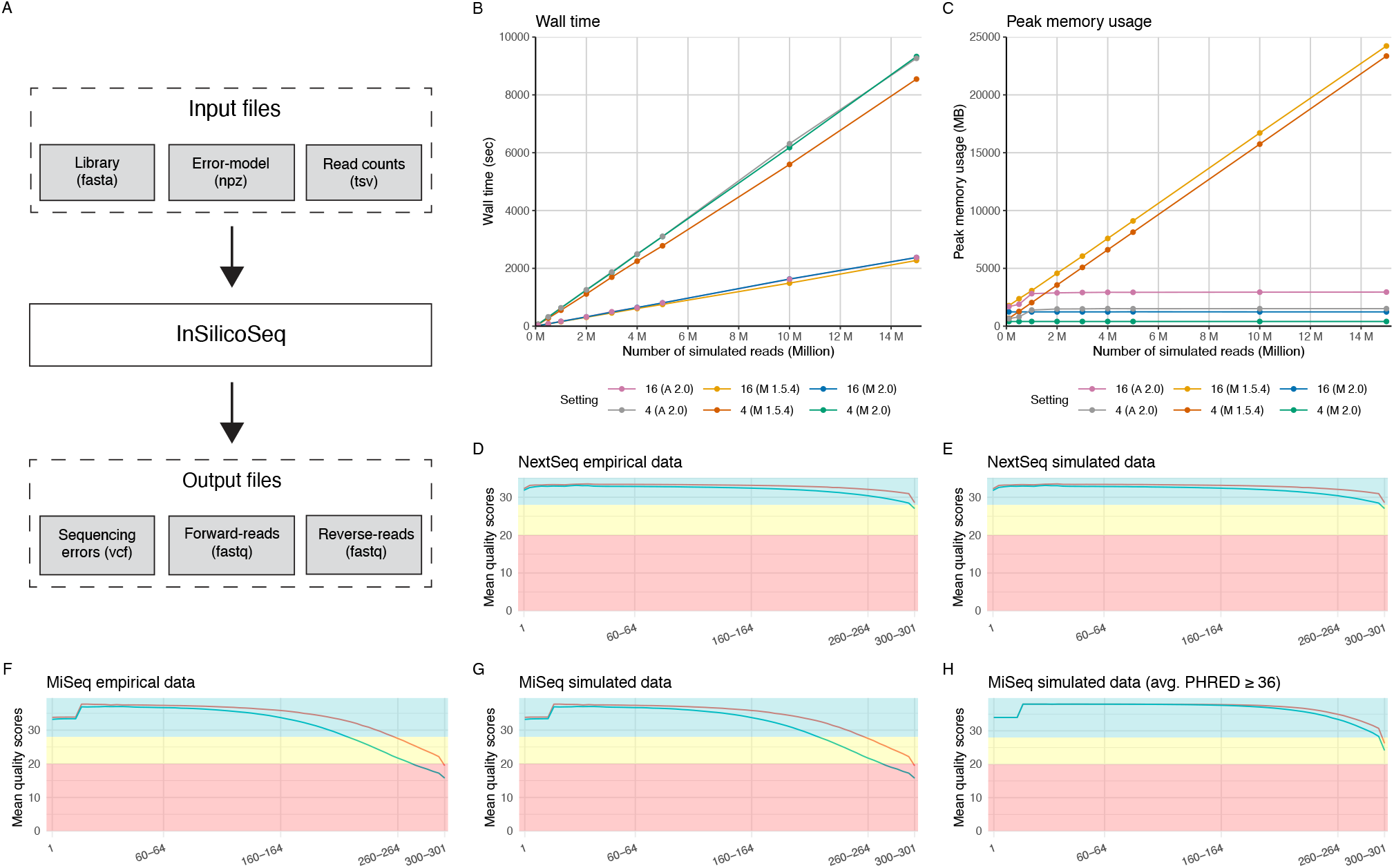
(**A**) Overview of the workflow to generate amplicon-based reads with InSilicoSeq 2.0. Benchmarking (**B**) Wall time and (**C**) Peak memory usage for amplicon and meta genome simulations executed by InSilicoSeq 2.0 and 1.5.4 with various number of reads performed on a cloud-based instance with 4 CPU’s (version 1.5.4 metagenome-mode in orange, version 2.0 metagenome-mode in green and amplicon-mode in grey) and 16 CPU’s (version 1.5.4 metagenome-mode in yellow and version 2.0 metagenome-mode in blue and amplicon-mode in pink). The Python module “memory_profiler” was used to determine the wall time and peak memory usage of each run^21^. Mean quality scores (Phred) per base of the forward (red line) and reverse (green line) 301base-pair reads based on: (**D**) empirical Illumina NextSeq data (Illumina base space project 370456276) and (**E**) simulations based on the empirical NextSeq data. Idem for (**F**) Empirical Illumina MiSeq data of SRA project PRJEB20178, (**G**) a simulation based on the empirical MiSeq data and (**H**) a simulation based on an error model of filtered empirical MiSeq reads with an average score of at least 36. The mean quality score plots have been generated using the “fastqcr” R-package.^19,20^

The type of sequencing data simulated by InSilicoSeq 2.0 can be controlled by the novel “-- sequence_type” parameter. The average fragment length of the paired-end reads can now be specified by the user via the “--fragment-length” and “--fragment-length-sd” parameters. The InSilicoSeq amplicon simulating mode requires a fasta-formatted file containing a library of amplicon sequences in combination with a read count file that contains, for each amplicon in the library file, the number of reads that will be generated (Figure 1A; example input files can be found as supplementary data 1-3). Finally, a precomputed error model can be selected to start the amplicon read simulation (Supplementary table 1). In addition, InSilicoSeq 2.0 provides the flexibility for the user to generate custom error models from BAM-files. The output of InSilicoSeq 2.0 contains the forward (R1), and reverse (R2) reads as separate fastq-files and for each read the sequencing errors are stored in a separate vcf-formatted file (Figure 1A; supplementary data 4) with the purpose of being used as ground-truth data for detecting and correcting sequencing errors.

### 2.2 Usability

An intra-version comparison was performed between the metagenome simulations generated by InSilicoSeq 1.5.4 and InSilicoSeq 2.0, and the amplicon-based simulations of InSilicoSeq 2.0. The aim of this comparison is to demonstrate the relevance of the improvements made in terms of useability and methodology of InSilicoSeq 2.0, in particular regarding issues that were shown to be problematic in the past^16 17^. The amplicon simulations of InSilicoSeq 2.0 were based on a library file that mimics an AIRR sequencing experiment based on Takara’s SMARTer Human BCR kit-like (Takara Bio USA) amplicons. The meta genome simulations of InSilicoSeq 1.5.4 and InSilicoSeq 2.0 were based on a referenced assembly for Baker’s yeast strain s288C (accession number: GCA_000146045). Both amplicon and metagenome simulations were based on simulating paired-end Illumina MiSeq based reads with lengths of 301bp. The intra-version comparison of InSilicoSeq was carried out on a Scaleway PRO2-XS cloud-based instance using 4CPU’s and 16 GB of RAM and a larger PRO2- M with 16 CPU’s and 64GB of RAM^17^. To avoid memory issues on the PRO2-XS instance for the InSilicoSeq 1.5.4 based meta genome simulations, the larger and more costly PRO2-M instance had to be used and downgraded to use 4 CPUs, illustrating the previous issues with scalability of InSilicoSeq 1.5.4. InSilicoSeq 2.0 had no issues with performing large simulations on the smaller PRO2-XS instance.^16^A direct comparison of the average per-base quality-score (Phred) of the simulated AIRR dataset to empirical sequencing data revealed that the quality of the reads simulated by InSilicoSeq 2.0 closely resemble the empirical sequencing data (Figure 1D-H). In addition, the intra-version comparison revealed that InSilicoSeq 2.0 now simulates the correct number of reads with accurate fragment sizes that were shown to be problematic in a previous benchmark with InSilicoSeq 1.5.4^16^ (Figure 1B and 1C and supplementary table 3, supplementary figure 1). Furthermore, the benchmark demonstrated improved scalability with consistent memory usage, independent of the simulation size (Figure 1B and 1C, supplementary table 2 and supplementary figure 1).

Finally, based on the monetary costs of operating the cloud-based instances, it would require within the range of €4.13E^-05^-€0.00018 to simulate 1 million bases using InSilicoSeq 2.0 (Supplementary table 2). Considering the estimated expense of $0.006 to sequence one million bases^18^, simulating an equivalent number of bases would require at most 1/100^th^ of the cost advocating for evaluating experimental designs through simulations prior to actual sequencing.

### 2.3 Benchmark

To evaluate the overall performance of InSilicoSeq 2.0, we performed our own benchmark using the latest versions of the popular read simulators ART (v 2.5.8)^11^, DWGSIM (v0.1.15)^12^, Mason (v2.0.9-e165baf)^13^, NEAT (v3.3)^14^, Wgsim (v0.3.1-r13)^15^. In this benchmark, we focused on 1) the ability of each tool to simulate forward and reverse reads with realistic Phred quality scores, 2) compute time and memory usage, and 3) fragment length distribution.

For benchmark 1), we have conducted a direct comparison using the tool FastQC^19^ (parsed with the fastqcr^20^ R-library) of simulated reads to empirical reads generated on Illumina HiSeq, NovaSeq, MiSeq and NextSeq platforms. The Phred score based comparison highlighted the capability of InSilicoSeq Kernel Density Estimation implementation to capture the Phred scores from the empirical reads. Therefore, InSilicoSeq’s reads generated by the four precomputed error models are superior compared to the default and precomputed error models of the other tools (Supplementary Figure 3).

For benchmark 2), time and peak memory usage is evaluated. Using the profiling tool memory_profiler^21^, each tool is profiled while simulating a varying number of reads (1, 3, 5, 7, 10 million) and read lengths (126pb, 151bp and 301bp), all based on the Baker’s yeast strain assembly s288C. All simulations were performed on a single cloud-based PRO2-M^17^ instance with 16 CPUs and 64Gb RAM. If a tool had the possibility to use more than one CPU, all 16 CPUs were assigned (More details can be found in Supplementary table 4). For each of the three series varying the read length, InSilicoSeq 2.0 showed stable performance (Supplementary Figure 4). Peak memory usage is of InSilicoSeq 2.0. is constant, similar to the tool’s art, dwgsim and wgsim (Supplementary Figure 4), while the run time scales linear and is relatively low.

For benchmark 3), the fragment length and fragment length standard deviation are evaluated for all tools. The Phred-score computations were based on 10 million read simulation. The mapping of the read pairs to the Baker’s years assembly genome was done using BWA mem (0.7.17-r1198-dirty)^22^. Fragment lengths have been extracted using a custom pySam^15^ based script. As expected, all tools produce correct fragment lengths (Supplementary Figure 5).

## 3. Conclusion

InSilicoSeq 2.0 has been designed to offer user-friendliness to both bioinformaticians and wet-lab scientists and can be executed on a wide range of computing systems. We demonstrated the novel amplicon-sequencing feature of InSilicoSeq 2.0 by creating simulations of paired-end reads of Adaptive Immune Receptor Repertoire (AIRR) data with a length of 301 base-pairs. In addition, we have added the 301 base-pair paired error model based on Illumina NextSeq data, as well as error models based on Illumina MiSeq data, considering various averaged Phred-scores to simulate amplicon reads across several quality- classes. We have shown, based on the AIRR sequencing example and the overall benchmark, that the software package produces realistic amplicon-based sequencing reads that closely resembles the Phred-scores of empirical reads. The performance assessment of InSilicoSeq 2.0 showed that the performance shortcoming reported in a recent and published benchmark study have been addressed, resulting in a scalable tool with constant memory usage regardless of the simulation size. Furthermore, the use of cloud-based instances enabled to estimate the monetary expenses to simulate 1M bases at a cost of €4.13E-05-€0.00018. This makes InSilicoSeq 2.0 an inexpensive approach for validating sequencing experiments as well as bioinformatic workflows and allows scientists to make an informed decisions on what settings will give the desired results.

## Availability

The software-package is freely available on https://github.com/HadrienG/InSilicoSeq under the MIT licence.

## Acknowledgements

The computations were also partially enabled by resources provided by the National Academic Infrastructure for Supercomputing in Sweden (NAISS) and the Swedish National Infrastructure for Computing (SNIC) at UPPMAX partially funded by the Swedish Research Council through grant agreements no. 2022-06725 and no. 2018-05973.

## SUPPLEMENTARY FIGURES

**Supplementary Figure 1:**
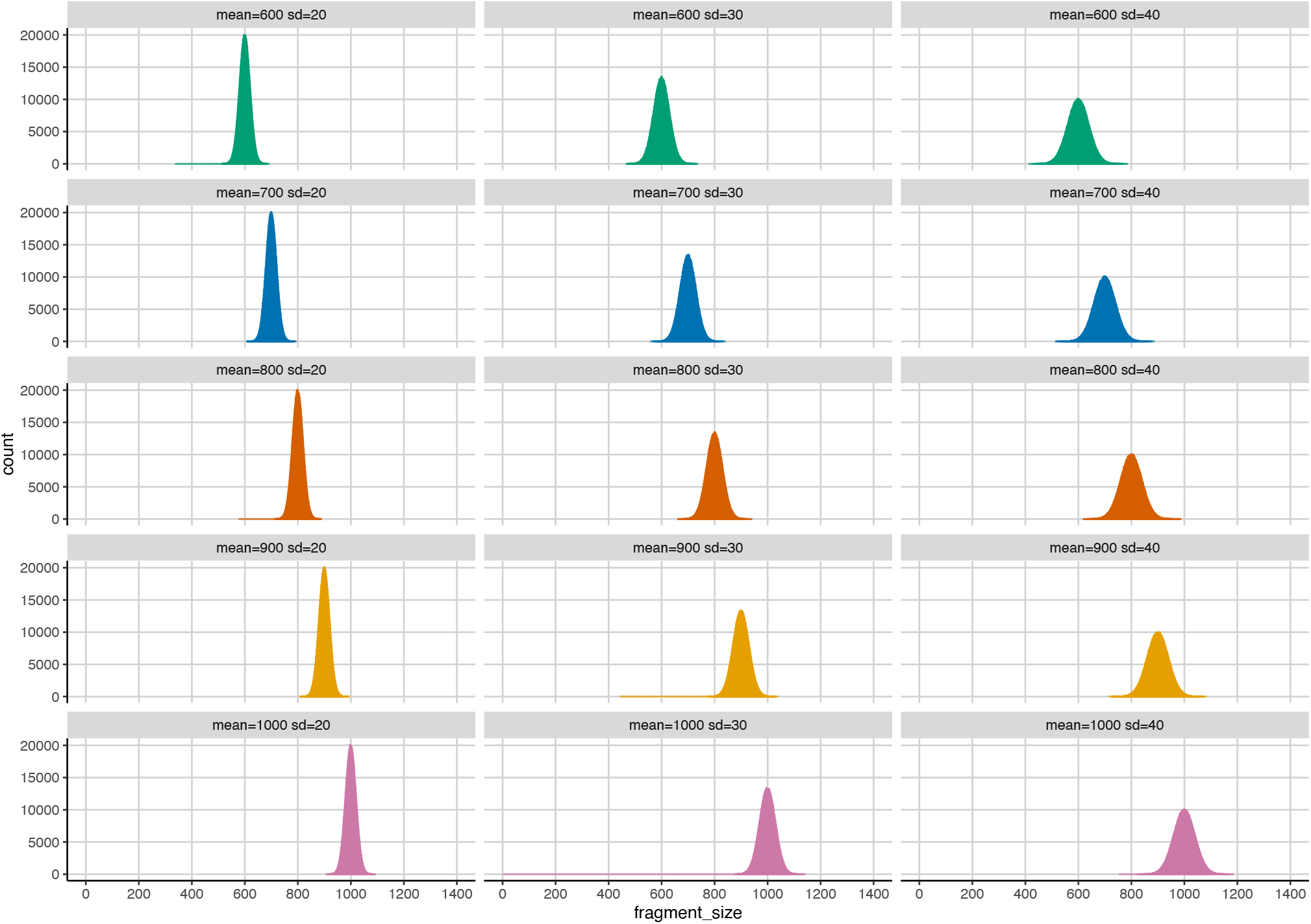
Overview of metagenome simulations based on the baker’s yeast genome with various fragment length. InSilicoSeq 2.0 have two new parameters to control the average length of the simulated fragments (--fragment-length) and the standard deviation (--fragment-length-sd). The bar plots show the distribution of fragment lengths for simulations with different average fragment length values. Results of simulations with an average fragment length of 600bp are shown in green, 700 bp in blue, 800 bp in red, 900bp in yellow and 1000 bp in pink. The columns show the result of simulations with different fragment length standard deviation values. Left column contains a standard deviation of 20bp, the middle column a standard deviation of 30bp and the right column a standard deviation of 40bp.

**Supplementary figure 2:**
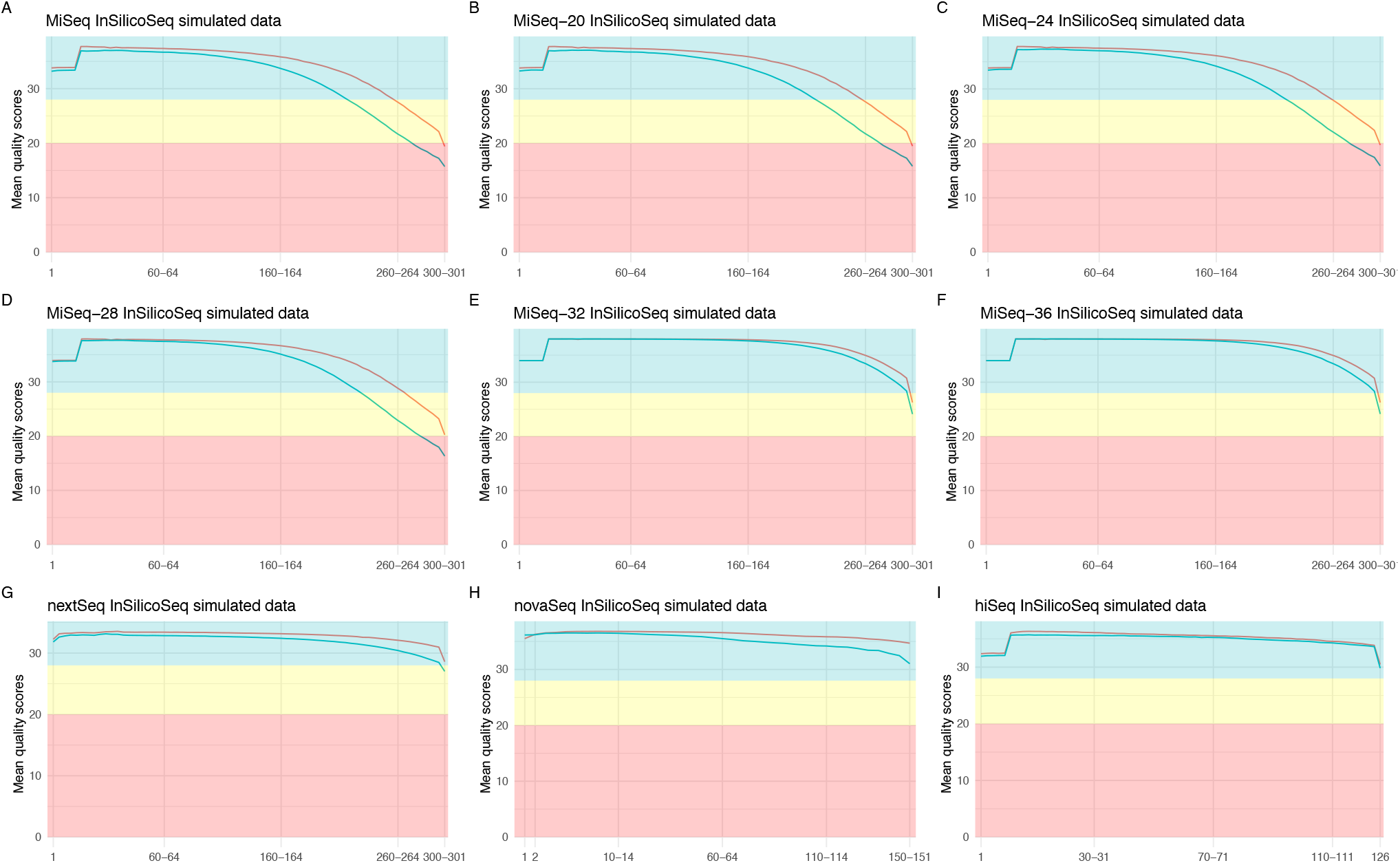
Overview of the per-base quality (Phred) scores for each precomputed error model that is available through the InSilicoSeq 2.0 cli. (**A**) default MiSeq error model with paired end reads of 301bp. MiSeq error models with paired end reads of 301bp and the forward and reverse reads have (**B**) an average quality score of at least 20 (**C**) an average quality score of at least 24 (**D**) an average quality score of at least 28 (**E**) an average quality score of at least 32 (**F**) an average quality score of at least 36. (**G**) NextSeq error model with paired end reads of 301bp (**H**) NovaSeq error model with paired end reads of 151bp (**I**) HiSeq error model with paired end reads of 126bp

**Supplementary Figure 3:**
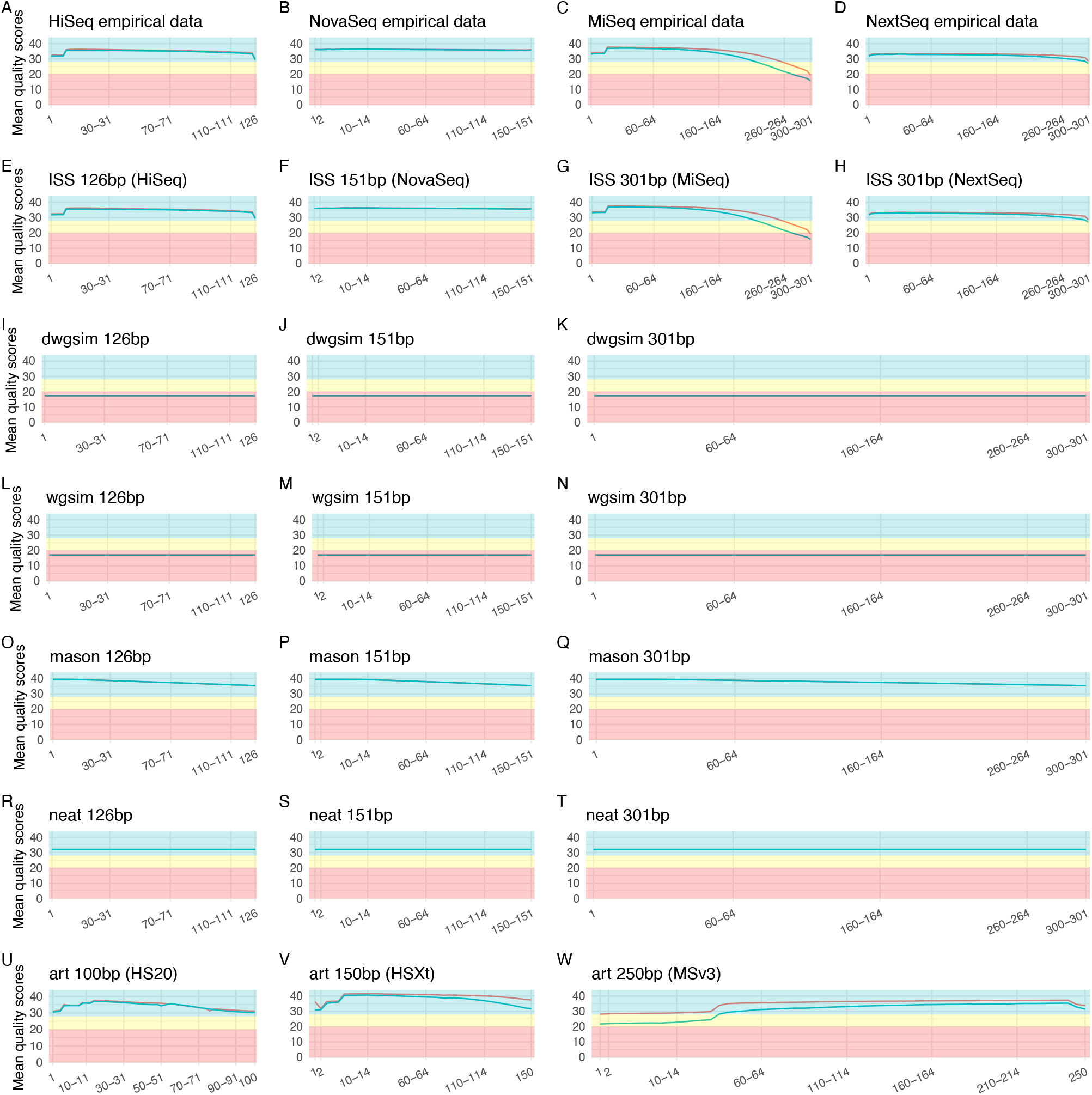
Benchmark of per base mean quality (Phred) scores of the forward (red line) and reverse (green line) reads of six tools compared to empirical data. The top row shows the per base mean quality (Phred) scores of empirical data: (A) HiSeq; paired 126bp reads (B) NovaSeq; paired 151bp reads (C) MiSeq; paired 301bp reads and (D) NextSeq; paired 301bp reads. The second row contains the per base mean quality (Phred) of the InSilicoSeq v2.0 precomputed (E) HiSeq (F) NovaSeq (G) MiSeq and (H) NextSeq based error models. Dwgsim, Wgsim, Mason, and Neat do not contain precomputed error models that can be used out of the box. For these tools the default error models were used with different read lengths (126bp, 151bp and 301bp) (L-T). For Art precomputed error models were used: (U) HS20: generated 100bp paired-end reads based on a HiSeq 2000 platform. (V) HSXt: HiSeqX plaporm with 150bp reads (No precomputed NovaSeq based error model is available). (W) MSv3: MiSeq v3 with 250bp reads

**Supplementary Figure 4:**
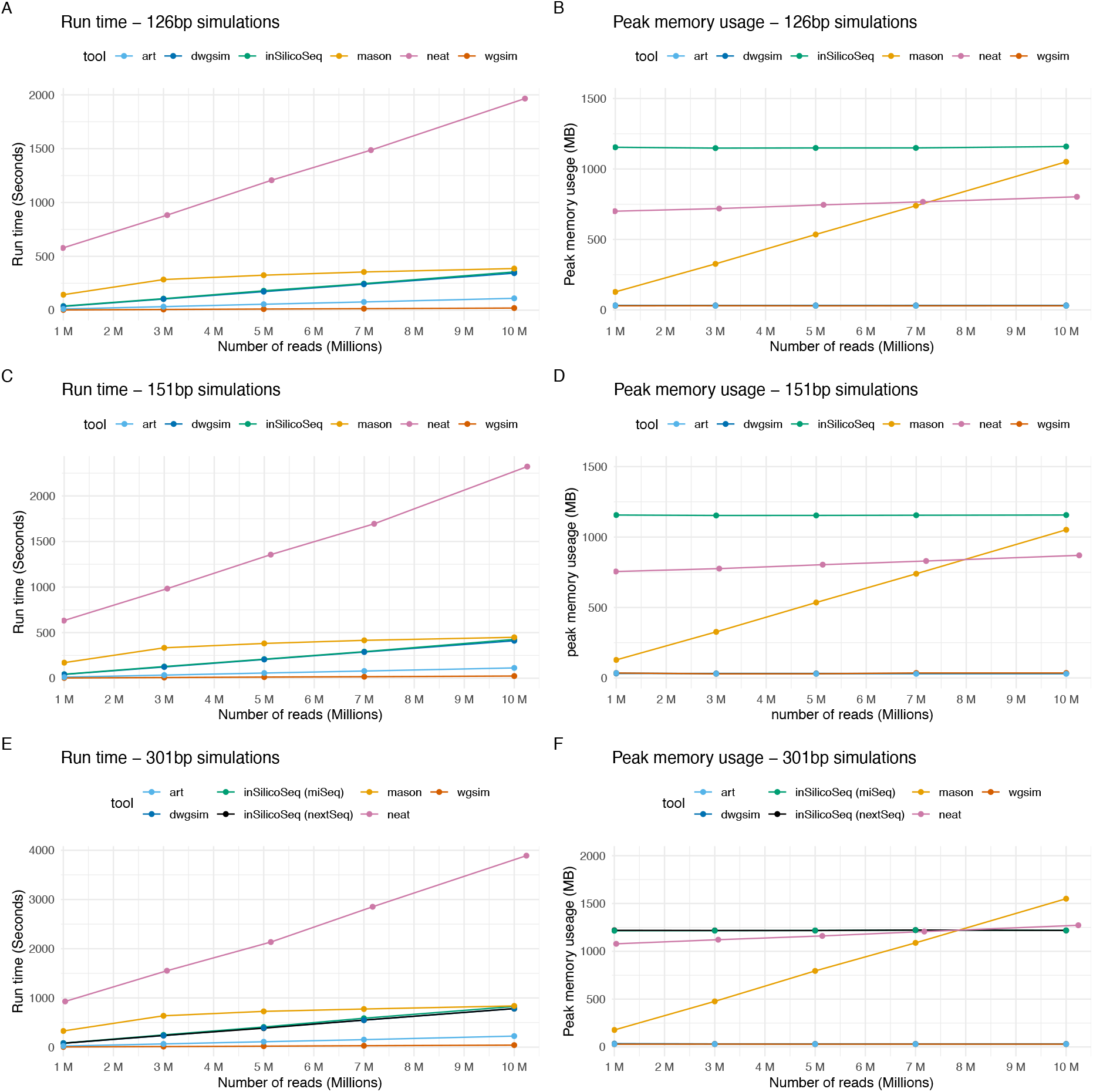
Benchmark of run time and max peak memory of the tools Art (light blue), Dwgsim (dark blue), InSilicoSeq (green and black), Mason (yellow), Neat (pink), and Wgsim (red) for simulations of 1,3,5,7 and 10 million reads. Simulations are based on the assembly for Baker’s yeast strain s288C. (A) Run time comparison and (B) peak memory usage of the 126bp simulations. None of the precomputed Art error models generated 126bp reads; therefore, the HS20 model was used generating 100 bp reads. (C) Run time comparison and (D) peak memory usage of the 151bp simulations. (E) Run time comparison and (F) peak memory usage of the 301bp simulations. InSilicoSeq have two precomputed models, MiSeq (green) and NextSeq (black), generating 301bp reads. Simulations from both models are included. Of general note: For Neat, the total coverage of the genome can only be specified instead of the total number of generated reads. In addition, this benchmark was performed on a cloud-based PRO2-M instance with 16 CPUs and 64Gb RAM. If the tools had possibility to specify the number of CPUs to use all 16 CPUs were assigned. Art (light blue) did not contain precomputed error models to generate paired end reads of length 126bp, 151bp and 301bp. Therefore, simulated data consisting of 100bp (HS20), 150bp (HSXt) and 250bp (MSv3) paired end reads were included instead. More details can be found in **Supplementary table 4**.

**Supplementary Figure 5:**
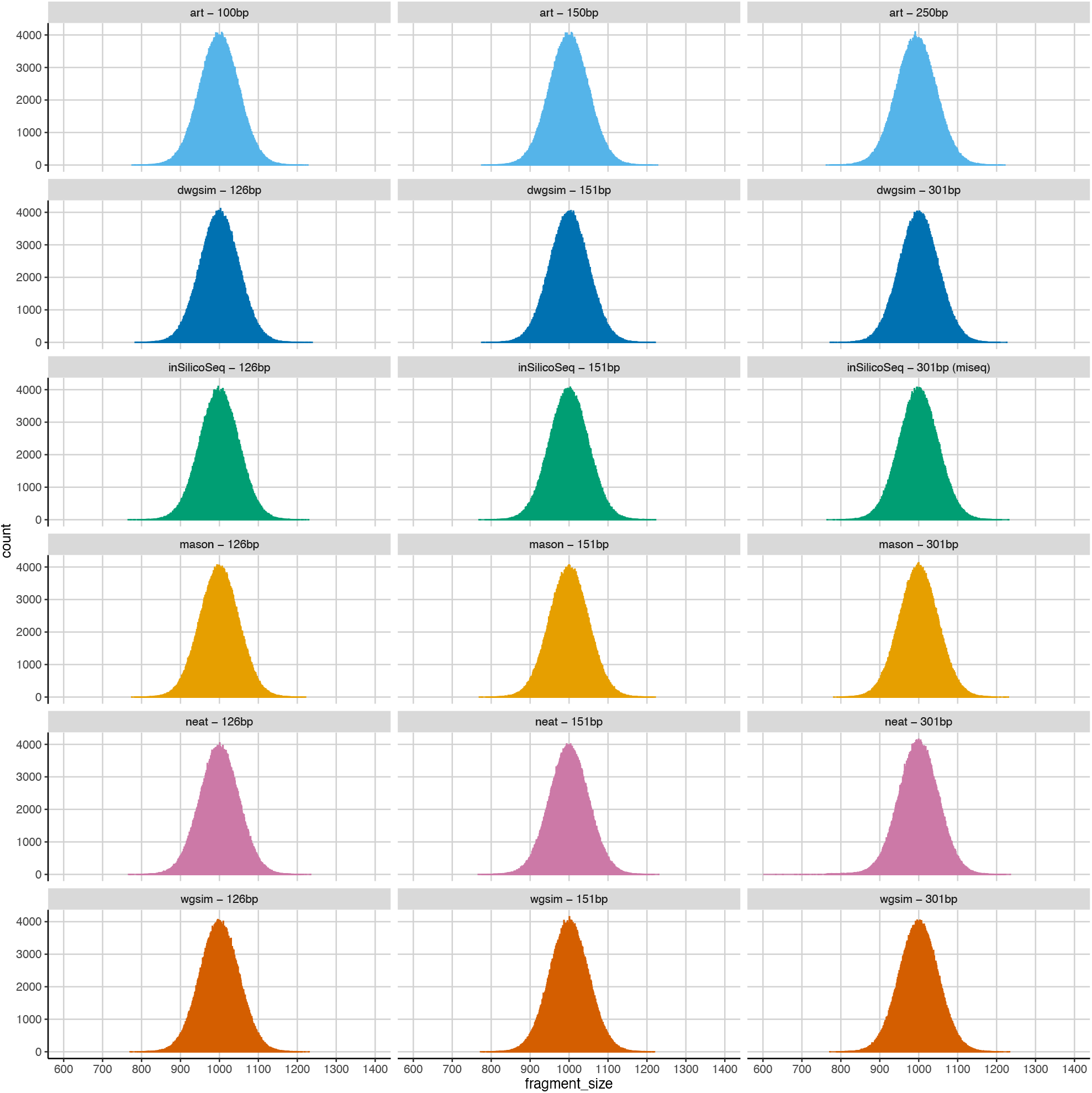
Benchmark of generated fragment length and fragment length standard deviations of Art (light blue), Dwgsim (dark blue), InSilicoSeq (green), Mason (yellow), Neat (pink) and Wgsim (red). Analyses are based on 1 million reads of 126bp (left column), 151bp (middle column) and 301bp (right column) long paired end reads based on the Baker’s yeast strain s288C and mapped to that reference using BWA mem^22^. Fragment lengths have been extracted using a custom pySam based script. For each simulation we have used a fragment length of 1000bp with a standard deviation of 50bp. Art (light blue) did not contain precomputed error models to generate paired end reads of length 126bp, 151bp and 301bp. Therefore, simulated data consisting of 100bp (HS20), 150bp (HSXt) and 250bp (MSv3) paired end reads were included instead.

## SUPPLEMENTARY TABLES

**Supplementary table 1:** Overview of the precomputed error models for different Illumina sequencing platforms that are available through the InSilicoSeq 2.0 cli.

| Error model | Sequencing platform | Read length | Filters | Cli |
| --- | --- | --- | --- | --- |
| nextSeq.npz | NextSeq | 2x 301bp | None | nextseq |
| miSeq_20.npz | MiSeq | 2x 301bp | Average Phred score $\geq 20$ | miseq-20 |
| miSeq_24.npz | MiSeq | 2x 301bp | Average Phred score $\geq 24$ | miseq-24 |
| miSeq_28.npz | MiSeq | 2x 301bp | Average Phred score $\geq 28$ | miseq-28 |
| miSeq_32.npz | MiSeq | 2x 301bp | Average Phred score $\geq 32$ | miseq-32 |
| miSeq_36.npz | MiSeq | 2x 301bp | Average Phred score $\geq 36$ | miseq-36 |
| miSeq.npz | MiSeq | 2x 301bp | None | miseq |
| hiSeq.npz | HiSeq | 2x125bp | None | hiseq |
| novaSeq.npz | NovaSeq | 2x150bp | None | novaseq |

**Supplementary_table_2.xlsx:** Performance benchmark of InSilicoSeq simulating AIRR sequencing based experiments with 301bp paired end reads. Simulations with 4 CPUs were performed on a PRO2-XS Scaleway instance (4 cores, 16Gb RAM, 100Gb block SSD, Ubuntu 22.04, with an estimated cost of €0.1258/Hour). Simulations with 16 CPUs were performed on a PRO2-M Scaleway instance (16 cores, 64Gb RAM, 100Gb block SSD, Ubuntu 22.04, with an estimated cost of €0.4538/Hour)

**Supplementary_table_3.xlsx:** Comparison of the expected number of reads and the number of simulated reads.

**Supplementary_table_4.xlsx**: Details of the simulated benchmark datasets generated by Art, Dwgsim, Wgsim, Neat, Mason and InSilicoSeq.

## SUPPLEMENTARY DATA-FILES

**Supplementary File 1: library.fasta**

**Supplementary File 2: read_count.tsv**

**Supplementary File 3: miseq_10K_reads.vcf**

**Supplementary File 4: miseq_10K_reads_R1.fastq**

**Supplementary File 5: miseq_10K_reads_R2.fastq**

**Supplementary File 6: run_iss.sh**

## References

1. Methé, B. A. et al. A framework for human microbiome research. Nature 486, 215–221 (2012).

2. Schoch, C. L. et al. Nuclear ribosomal internal transcribed spacer (ITS) region as a universal DNA barcode marker for Fungi. Proc. Natl. Acad. Sci. 109, 6241–6246 (2012).

3. Robins, H. S. et al. Comprehensive assessment of T-cell receptor β-chain diversity in αβ T cells. Blood 114, 4099–4107 (2009).

4. Wu, X. et al. Focused Evolution of HIV-1 Neutralizing Antibodies Revealed by Structures and Deep Sequencing. Science 333, 1593–1602 (2011).

5. Lind, A. L. & Pollard, K. S. Accurate and sensitive detection of microbial eukaryotes from whole metagenome shotgun sequencing. Microbiome 9, 58 (2021).

6. Andreu-Sánchez, S. et al. A Benchmark of Genetic Variant Calling Pipelines Using Metagenomic Short-Read Sequencing. Front. Genet. 12, 648229 (2021).

7. Ewing, B. & Green, P. Base-Calling of Automated Sequencer Traces Using Phred. II. Error Probabilities. Genome Res. 8, 186–194 (1998).

8. Gourlé, H., Karlsson-Lindsjö, O., Hayer, J. & Bongcam-Rudloff, E. Simulating Illumina metagenomic data with InSilicoSeq. Bioinformatics 35, 521–522 (2018).

9. Lantbruksuniversitet, S. PRJEB20178. https://www.ncbi.nlm.nih.gov/bioproject/PRJEB20178 (2018).

10. Illumina base space demo files. https://basespace.illumina.com/datacentral.

11. Huang, W., Li, L., Myers, J. R. & Marth, G. T. ART: a next-generation sequencing read simulator. Bioinformatics 28, 593–594 (2012).

12. DWGSIM. https://github.com/nh13/DWGSIM.

13. Holtgrewe, M. Mason: a read simulator for second-generation sequencing data. Technical Report FU Berlin (2011).

14. Stephens, Z. D. et al. Simulating Next-Generation Sequencing Datasets from Empirical Mutation and Sequencing Models. PLoS ONE 11, e0167047 (2016).

15. Li, H. et al. The Sequence Alignment/Map format and SAMtools. Bioinformatics 25, 2078–2079 (2009).

16. Milhaven, M. & Pfeifer, S. P. Performance evaluation of six popular short-read simulators. Heredity 130, 55–63 (2023).

17. Scaleway Instance pricing. https://www.scaleway.com/en/pricing/?tags=compute.

18. Wetterstrand, K. A. DNA Sequencing Costs: Data from the NHGRI Genome Sequencing Program (GSP). https://www.genome.gov/sequencingcostsdata.

19. Andrews, S. A quality control tool for high throughput sequence data. https://www.bioinformatics.babraham.ac.uk/projects/fastqc/ (2010).

20. Kassambara, A. fastqcr: Quality Control of Sequencing Data. https://rpkgs.datanovia.com/fastqcr/index.html (2023).

21. Pedregosa, F. & Gervais, P. memory profiler. https://github.com/pythonprofilers/memory_profiler.

22. Li, H. Aligning sequence reads, clone sequences and assembly contigs with BWA-MEM.

