## Supplementary material for "InSilicoSeq 2.0: Simulating realistic amplicon-based sequence reads": run_iss.sh

```
#install InSilicoSeq
pip install InSilicoSeq

# run InSilicoSeq
iss generate \
--sequence_type amplicon \
--model "miseq" \
--genomes "library.fasta" \
--readcount_file "read_counts.tsv" \
--store_mutations \
--output "${PWD}/miseq_10K_reads" \
--cpus 4 
```
